# Two closed systems for long-term propagation of the marine tunicate Botryllus schlosseri isolated from natural seawater

**DOI:** 10.1101/2025.07.01.662663

**Authors:** Jens Hamar, Weizhen Dong, Brenda Luu, Mandy Lin, Isabel Enriquez, Maxime Leprêtre, Alison M. Gardell, Baruch Rinkevich, Dietmar Kültz

## Abstract

Advanced methodologies for Botryllus schlosseri artificial seawater systems are needed to decrease dependency of large-scale culture on natural seawater and expand use of this important new model organism to more inland laboratories. We constructed two botryllid tunicate customized closed aquaculture systems, a static system consisting of lightly aerated jars fed with commercial filter feeder diet, and a recirculating aquaculture system (RAS) consisting of standard marine RAS components fed live microalgae and zooplankton diets. Initially, static tunicate culture yielded exponential growth in contrast to the RAS system, which yielded poor survival and negligible growth. Modifications were made to the RAS system to improve water treatment proficiency that greatly improved tunicate survival and growth. Experiments were performed isolating feed and water type as variables that differed between the static and RAS systems to evaluate their effects. A live feed combination achieved five-fold greater growth relative to a commercial concentrate diet. B. schlosseri maintained in optimized RAS water achieved two-fold faster growth relative to animals maintained with freshly prepared artificial seawater indicating that the RAS water was beneficial to the animals. Feeding frequency of the RAS system was increased from three times per week to daily. The RAS system and procedural modifications resulted in comparable growth rates in the static and RAS systems. Both optimized systems are suitable for long-term propagation and sustenance of botryllid tunicate populations supporting both sexual and asexual modes of reproduction with a current residence time of over 24 months.

## 1. Introduction

Many of the scientific advances relevant to human health and nutrition were initiated in simplified non-human organisms. Notable examples include: the discovery of gene arrangement and genetic inheritance through chromosomes using Drosophila[1], the molecular basis of vertebrate development using zebrafish[2], and initial characterization of human cyclins in yeast[3]. Many more advances in mechanistic characterization of biological processes are needed to address modern problems associated with human health and disease. For example, understanding immune recognition pathways and tissue or organ regeneration processes requires further development of suitable model organisms to reveal better insight into the underlying mechanisms. The emergence of tunicates as invertebrate models of vertebrate physiology is supported by their status as an ideal clade of organisms that has many unique physiological abilities (e.g. whole body regeneration). This model also combines favorable attributes of invertebrate models in terms of their easy and flexible reproduction strategies and the ability to clone new animals while retaining high relevance for vertebrate physiology due to their close phylogenetic proximity as a chordate[4]. More specifically, the colonial tunicate B. schlosseri has already demonstrated high value as a model for many important biological processes including allorecognition[5], aging, stem cell biology[6], regeneration[7], and ecotoxicity[8]. Routine use of these animals in scientific studies requires immediate access to wild populations or, in most cases, clean and genetically defined strains of laboratory populations that can be easily propagated under reproducible, stable conditions. Tunicate propagation is typically done using seawater directly sourced from coastal waters. Methods of static B. schlosseri aquaculture have been established involving regular water changed with natural seawater[9], but practical application of these approaches require marine laboratories with convenient coastal access. Advancing knowledge and expertise in proficient aquaculture of these animals using artificial seawater systems would make this powerful new model accessible to a vastly larger range of laboratories.

An alternative to naturally sourced seawater is to prepare it from dehydrated salt mixes. Such synthetic seawater is commonly used to maintain marine animals in closed systems. However, in flow through or direct replacement systems, synthetic seawater is expensive, ecologically unsustainable, and impractical on a large scale. Recirculating aquaculture systems (RAS) have been identified as a means to minimize cost by reduced use of consumable resources (i.e. water and sea salt) by conditioning and reuse of water[10]. They can create artificial marine environments that are ecologically sustainable and suitable for the long-term culture of marine species in areas where natural seawater is unavailable, or its use is impractical. Additional benefits include better control over environmental/abiotic parameters[11], conservation of invested cost into modification of the source water (such as temperature change or salinity adjustment), increased biosecurity through protection from potentially harmful impacts via the source water (i.e. introduction of toxins and pathogens), and reduction of waste effluent to the environment.

In many areas recirculating aquaculture systems (RAS) have made aquatic culture possible, or at least more cost effective, where suitable water sources are limited[12] and allowed for inland culture of diverse phyla of marine species including fish (i.e. clownfish[13,14], pompano[15]) and invertebrates (crustaceans, echinoderms[16], molluscs[17], and corals[18]) in inland areas without reliance on natural seawater. A proficient RAS has recently been established for another colonial tunicate (Botrylloides diegensis)[19]. The objective of this project is to build on this prior advancement and eliminate reliance on natural seawater for the culture of B. schlosseri by constructing artificial seawater systems (both static and RAS) that are optimized for generating a steady supply of healthy, actively growing animals that can robustly support scientific research using this intriguing model organism.

## 2. Materials and Methods

### 2.1 Animals

Parent stock animals were collected from Northern California floating docks within Berkely Marina (BM), Bodega Bay (BB), and San Francisco Harbor (SFH). Adult wild collected B. schlosseri and Botrylloides spp. were collected and held jointly in the initial RAS system only after the first field collection. Wild-caught animals from all other field collections were brought into the laboratory and held in isolation tanks for collection of B. schlosseri larvae. Larvae were collected onto 2” X 3” glass slides from aggregates of adult animals approximately 1 inch in diameter using two methods: (1) By tying colonies to slides and placing them across from a clean slide in a holder; and (2) By placing the aggregate within a box with glass slides lining the interior. Clean slides colonized by the free-swimming larval stage (1 to 3 days post collection of wild colonies) were removed and transferred to animal housing units (AHU). Slides were examined using a Leica EZ4W microscope every 3 to 5 days. Colonies showing heavy contamination, poor health, or lack of growth were culled from the slide. To decrease the density of slides containing multiple healthy colonies, single colonies of 4 to 6 zooids were transferred to new slides using a razor blade as previously described[20].

### 2.2 Animal Housing

Long term growth and propagation of selected colonies was achieved in AHUs specific to the culture system. AHUs of the static system consisted of lightly aerated 2.8 L static glass jars. AHUs of the RAS system consisted of one 115 L glass aquarium and flow through clear plastic zebrafish tanks from Aquaneering Inc: four 9.5 L tanks (cat# ZT950), and fourteen 1.8 L tanks (cat# ZT180). Static system slides were held in glass slide holders. To reduce algae growth underneath the tunic caused by light penetrating the back side of slides, the RAS system slides were held in custom manufactured slide holders with opaque black acrylic backing. Ten slide capacity holders were used for the 9.5 L AHUs and a single slide capacity holder for the 1.8 L AHUs.

### 2.3 Feeding and Maintenance

Initially, tunicates in both systems were fed a mixture of Roti-Rich (Florida Aqua Farms Inc) and Phyto-Feast Live (Reed Mariculture Inc). Subsequently, RAS system tunicates were transitioned to a live feed mix consisting of three microalgae species (Tetraselmis spp, Nannochlorpsis spp, and Isochrysis galbana) and marine rotifers (Brachionus plicatilis). Animals cultured in the Static system were fed twice per week while RAS system tunicates were initially fed every other day. The revised RAS system feeding frequency was eventually increased to daily to increase growth rates. For both systems, artificial seawater (AS) was prepared using a dry artificial seawater mix (Red Sea Coral Pro Salt, product# R11065). For the static system, 100% water changes were performed using 30 ppt AS once per week on each AHU. For the RAS system, 24 L of water was syphoned from the refugium (RFG) and replaced with fresh AS. Salinity and temperature for all systems were maintained at 30 ppt and between 19 and 20°C, respectively. Colonies were cleaned weekly using soft brushes. Debris surrounding the tunicate colonies was carefully removed by scraping the glass slides, taking care not to damage the colony ampullae. Ammonia, nitrite, and nitrate were checked periodically using API test kits (product numbers LR8600, 26, and LR1800 respectively).

### 2.4 Live Feed Culture

All microalgae strains were cultured in 2 L glass jars with flat glass top and a bottom drain with a dispensing valve. Vessels were exposed to continuous lighting using a 37 watt LED light (Finnex StingRAY 2 - 48" LED Aquarium Light, model#LT-FX-FL48) and lightly aerated with air filtered through a 0.22 micron syringe filter. Each vessel received 1 L of sterile 33 ppt AS supplemented with 200 µl part A and 200 µl part B of Fritz Aquatics F2 media per L AS twice per week. Rotifers were maintained in an aerated plastic 8 L conical bottom container with daily 10% water changes and feeding of resuspend dry yeast powder (Saccharomyces cerevisiae).

### 2.5 Initial RAS System

The RAS system was constructed using a 124 cm H X 168 cm L X 56 cm W stainless steel rack as the frame and including the following standard marine husbandry components: a 30 cm H X 132 cm L X 79 cm W, 150 L plastic stock tank (Rubbermaid mfg #424300BLA) as the sump (SMP); a circulating water pump (PMP), 20 gpm submersible pump (Danner model#12B); a biofilter (BFR) consisting of a 89 cm H X 58 cm diameter 210 L polyethylene drum filled with 30 L of 3.8 cm diameter plastic media bioballs (Bulk Reef Supply product# 000641); a 20” mechanical filter (MFR) containing a 200 µM cartridge filter; two temperature control chillers (CHL), AquaEuro Max Chill 1/13 HP chiller (model# MC-1/13HP); a refugium (RFG) contained in a 30 cm H X 69 cm L X 51 cm W 115 L glass tank illuminated by a 26 watt full spectrum LED light (Tunze cat #8850); a foam fractionator (FFR), Bubble Magus Curve 9 (model # BM-CURVE 9); and a 40 watt UV sterilizer (UVS), IceCap (product# IC-HOUV-40W). Pressurized water lines from the PMP and gravity feed return lines were constructed from 3/4” and 1.5” schedule 40 PVC pipe, respectively.

### 2.6 Static system

The static system was modeled after that previously described in Rinkevich and Shapira[20]. Larvae of wild colonies were spawned and attached to glass slides at the UC Davis Cole B facility within one week after field collection. These lab-born colonies developed from oozooids were maintained vertically on glass slides in 2.8 L glass tanks containing 30ppt standing AS at a constant temperature of 201°C and aerated by air stones as previously described[20]. All genotypes used in this study were reared in individual tanks to avoid competition[21]. AS for each tank was changed every week and colonies were gently cleaned once a week using soft brushes. All experimental colonies were in good health and have been born and reared in stable lab conditions for at least 3 months.

### 2.7 Revised RAS System

The revised RAS system includes all the components from the initial system with the following modifications: Total system circulation was divided into two water pumps (PMP), PMP1 = 20 gpm submersible pump (Danner model#12B) and PMP2 = variable speed DC submersible pump (Simplicity 2100DC Pump, model# SIM2005); a 38 L foam fractionator tank (FFT) was added to contain the FFR, 10 Gallon Polyethylene Tamco® Tank - 13" Dia. x 19" Hgt.( item # 3005); and a 64 L polyethylene mixing tank (MXT) was added to contain PMP2, 17 Gallon Polyethylene Tamco® Tank - 18" Dia. x 15" Hgt. (item # 3010).

### 2.8 Growth quantification

Growth data were collected for individual genotypes defined as all asexually produced progeny from a single sexually produced oozooid. For each recorded genotype, the numbers of individual animal units of a colony (zooids) were periodically recorded. The number of systems (a system being a discrete cluster of zooids averaging 8-12 zooids under a common tunic) was recorded as the unit of growth in the revised RAS system during a period of 3 weeks in 2025 when no animals were harvested for experiments.

### 2.9. Feeding Experiment

Two slides per genotype (average of 3 systems, 30 zooids per slide) from three static system raised tunicate genotypes were selected for exposure to two feeding treatments: The commercial feeding regime (CF) described above for the static culture system and the live feed regime (LF) used for the RAS system. Each treatment group received one slide from each genotype for a total of six replicates per treatment. All replicates were cultured as described above using the static system in 2.8 L jars with light aeration. Protein concentration (quantified for each individual feed component using a BCA assay (Thermo-Pierce) was used to adjust feed component volumes to normalize feed quantity between treatment groups. For the commercial feed group, 1 ml of Roti-Rich and 1 ml of Phyto-Feast was mixed into 10 ml of water, from which 1 ml was distributed to each replicate. For the live feed group, 57 ml of each microalgae culture was mixed with 225 ml of rotifer culture from which 150 ml was distributed to each replicate. Zooid counts and imaging was performed on each replicate at weekly intervals for 3 weeks. For each treatment group, mean relative change in zooid number (RCZN) was calculated as the total number of zooids at the final time point divided by the number of zooids at the start of the experiment for each genotype.

### 2.10. Water Quality Experiment

Ten individual genotype RAS system raised slides of tunicates (one 6-7 zooid system/slide) were randomly allocated to two treatment groups (5 replicates/group): One group received only freshly prepared AS and the other received water from loop 1 of the revised RAS system. With the exception of the water source for water changes, all replicates were cultured as described above using the static system in 700 ml glass jars with light aeration. Imaging and zooid counts was performed on each replicate twice per week for 3 weeks. For each treatment group, RCZN was calculated as described above.

### 2.11. Statistics

All statistical analyses were performed using Rstudio version 2021.09.1. One-tailed Welch and two sample t tests were performed on all RCZN comparisons.

## 3. Results

### 3.1. Initial RAS System

The completed 660 L total volume initial RAS system (Fig. 1A) was arranged with three elevated levels on the stainless steel rack: The top level supported 5 AHUs (one 115 L and four 10 L); the second level supported 14 1.8 L AHUs; and the bottom level supported the 115 L refugium and two CHLs. The SMP and BF were located at opposite ends of the rack. The SMP was the lowest point to which all water from the system flowed back to by gravity and contained the single circulation PMP and the FF. Pressure driven water from the PMP was split into two paths: One path directed flow through the MFR, then through the CHLs, and then to the RFG which gravity fed back to the SMP, and the second path directed flow through the UVS and to two distribution manifolds feeding the top and second level AHUs. Effluent water from the top-level AHUs gravity-fed into the BF and then back to the SMP. Effluent from the second level AHUs gravity-fed directly back to the SMP. The refugium was illuminated with the LED light mounted to the bottom of the second level and populated with macrophyte fragments and macrophyte-colonized rocks composed of five different species collected from the BBM collection site (Fig. 1B). In combination, the water turnover rate and assimilation of nitrogenous wastes by refugium macrophytes, were sufficient to keep system ammonia, nitrite, and nitrate below detectable limits.

**Figure 1.**
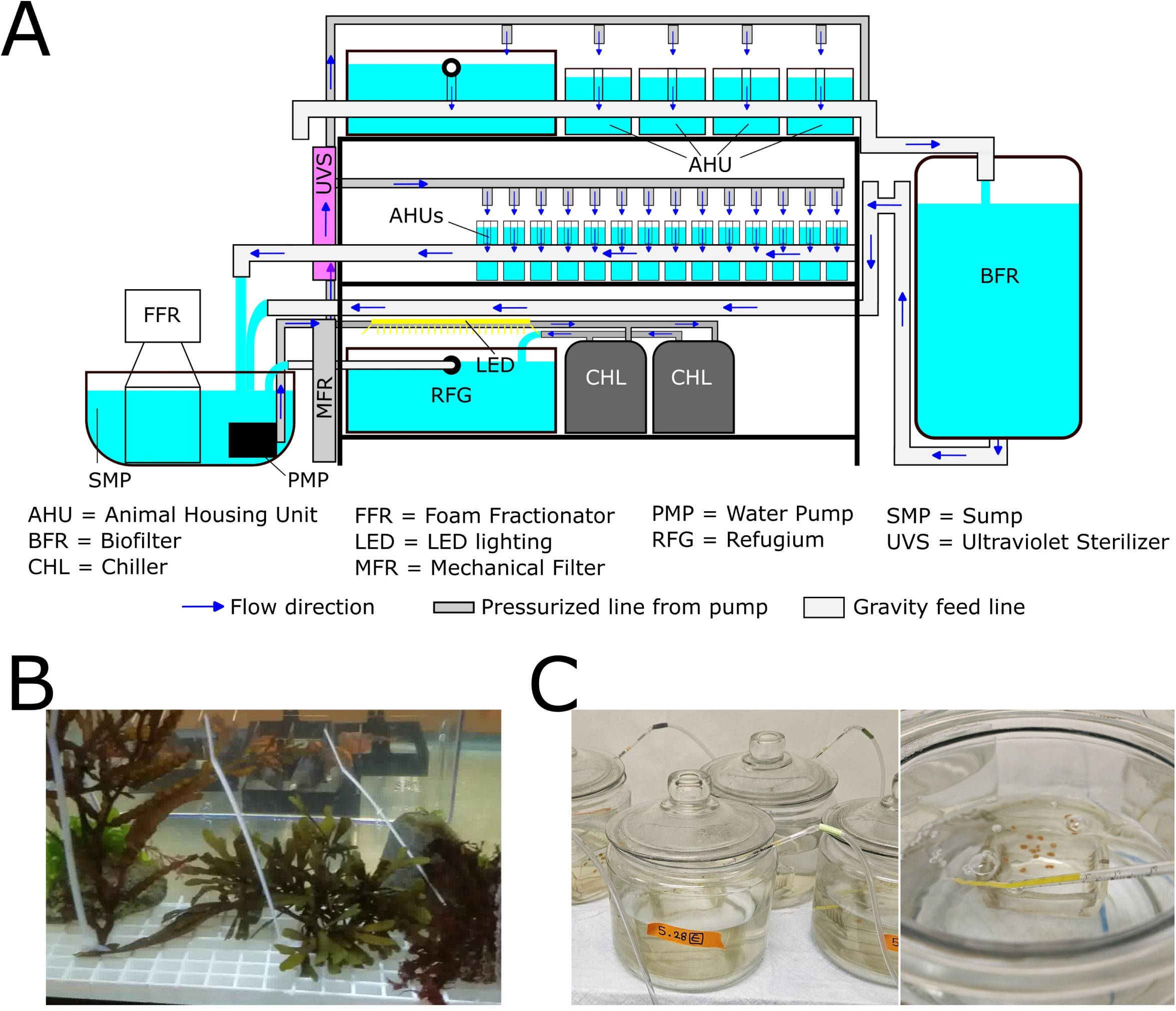
Initial culture systems. Schematic of initial RAS system with flow directionality (A) and macrophyte population of the refugium (B). Images of static system AHUs (C).

### 3.2. Static System

The completed static system consisted of 20 individual AHUs (Fig. 1C) each containing a single B. schlosseri genotype divided among three levels of a 142 cm H X 137 cm L X 33 cm W stainless steel rack. The single aerator distributed air to each AHU through individual lines connected to a filtered 1 ml serological pipette. The water turnover rate was sufficient to keep ammonia levels below detectable limits.

### 3.3. Growth rates in initial aquaculture systems

In the static system, tunicates rapidly expanded from single oozooids to multi system colonies (Fig. 2A), exhibiting exponential growth at a rate of approximately 0.46 per week, equivalent to a doubling time of 10.6 days, for selected genotypes (Fig. 2B). In the initial RAS system some oozooids grew to a maximum of six zooids with only a couple systems surviving up to 100 days (Fig. 2C). These surviving systems would eventually regress and disappear within another 60 days (Fig. 2D). There were no surviving oozooids from the first BM collection beyond 100 days. Only three and two oozooids obtained from BB and SFH collections, respectively, survived beyond 100 days. The initial RAS system was able to support Botrylloides spec. growth much better than growth of B. schlosseri. Easily noticeable continuous colony expansion was observed for Botrylloides spp. This qualitative visual observation was not substantiated with quantitative data as Botrylloides spp. was not the target species (Fig. 2E). Expansion of Botrylloides colonies continued for over a year in the initial RAS system until they were culled to eliminate competition with B. schlosseri.

**Figure 2.**
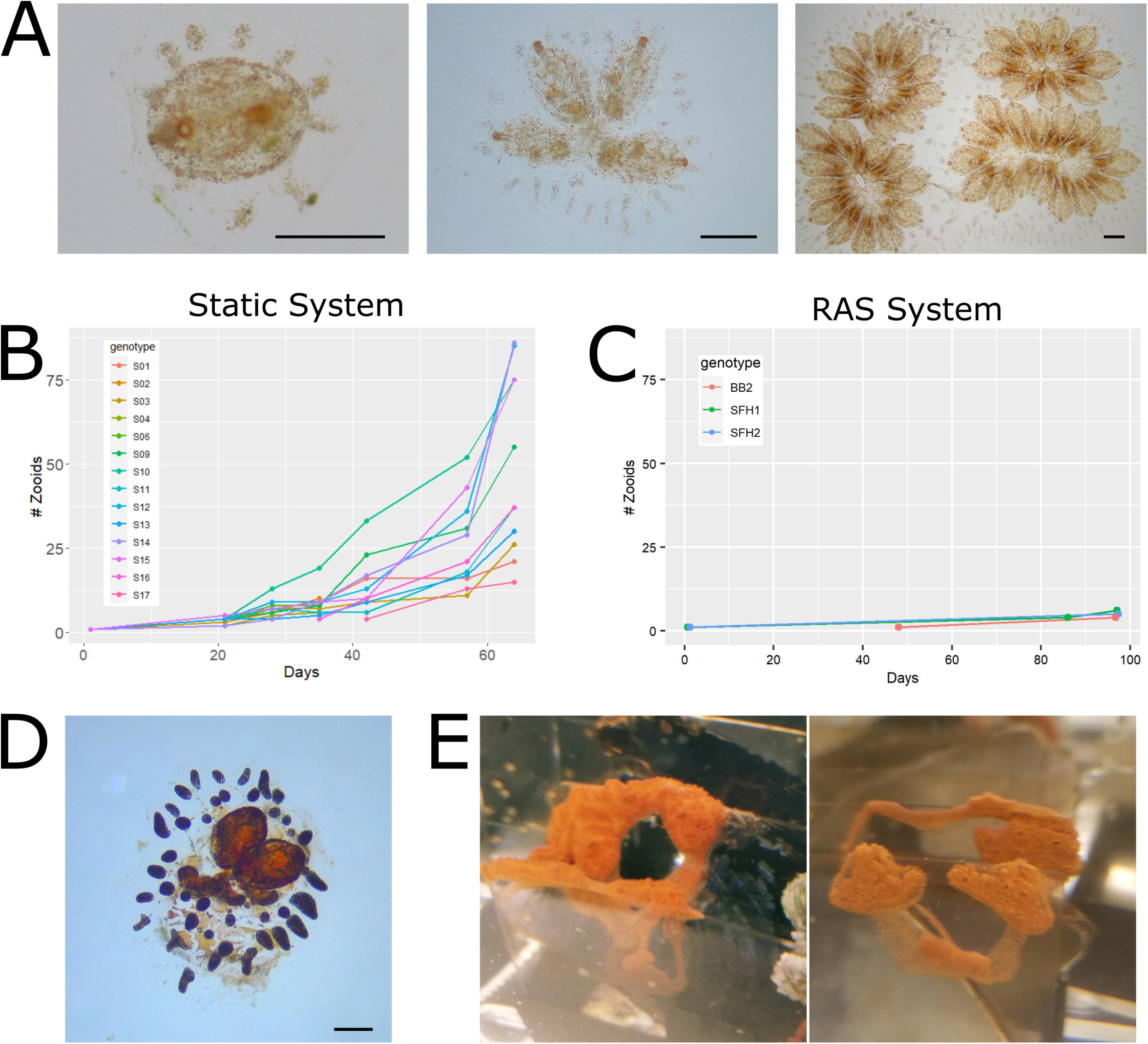
Tunicate growth in initial culture systems. Images of growth progression from a single oozooid to multi-zooid system to multi-system colony in static culture system (A). Growth of individual Botryllus genotypes quantified as number of zooids over days in culture of the static system (B) and the few surviving genotypes in the initial RAS system (C). Images of regressing Botryllus genotype (D) and expanding Botrylloides in initial RAS system. Scale bars in each micrograph = 1 mm.

### 3.4. Revised RAS system

Water circulation in the 760 L total volume newly optimized, revised RAS system (Fig. 3A&B) is divided into two independent loops driven by separate PMPs. Loop 1 starts in the lowest elevation point of the system (SMP) with PMP1 which drives water flow at a rate of 14 lpm through a MFR, the UVS, then diverges into two lines feeding the AHUs on level 2 and level 3. The effluent from both levels of AHUs collects into gravity-fed lines that flow directly into the FFT containing the FFR. The FFT overflow supplies water by gravity to the MXT. The resulting MXT overflow passes through the RFG illuminated by two LEDs run at half power, before returning back to the SMP. Loop 2 starts in the MXT with PMP2 driving water at a rate of 27 lpm into two lines: one flows directly into the BFR, and the other flows into another MFR, then through the two CHLs, and into the BFR. From here, water flows by gravity back into the MXT where interchange between the two loops occurs. Without input from loop 1, loop 2 circles back on itself between the BFR and the MXT. If loop 2 is off, loop 1 cycles between AHUs, tank 1, tank 2, the refugium, and the sump. This allows loop 1 to be turned off preventing immediate flushing of feed from the AHUs while feeding the animals. Relative to the initial RAS system, this arrangement allows maintenance of a high flow rate of loop 2 and supports better thermal management; increased mechanical and biological filtration efficiency; redundancy in the flexibility to operate the entire system with one loop in the case of an equipment failure; minimized flow through loop 1 which increases UV sterilization, longer retention time of feed in AHUs, and longer retention time of AHU effluent in FFR leading to increased efficiency of removing excess feed and deleterious tunicate waste. This new arrangement also allows animals to assimilate potential beneficial feed generated by the refugium before it is removed by the rest of the filter system. B. schlosseri colony-containing slides were held with either a single slide holder in 1.8 L AHUs of level 2 (Fig. 3 C&D) or ten slide holders in 9.5 L AHUs of level 1 (Fig. 3E). At the time of the RAS remodel, all but two macrophyte varieties, Gracilaria sp and Porphyra sp (Fig. 3F), had regressed and disappeared. These two remaining species grew rapidly and repopulated the refugium.

**Figure 3.**
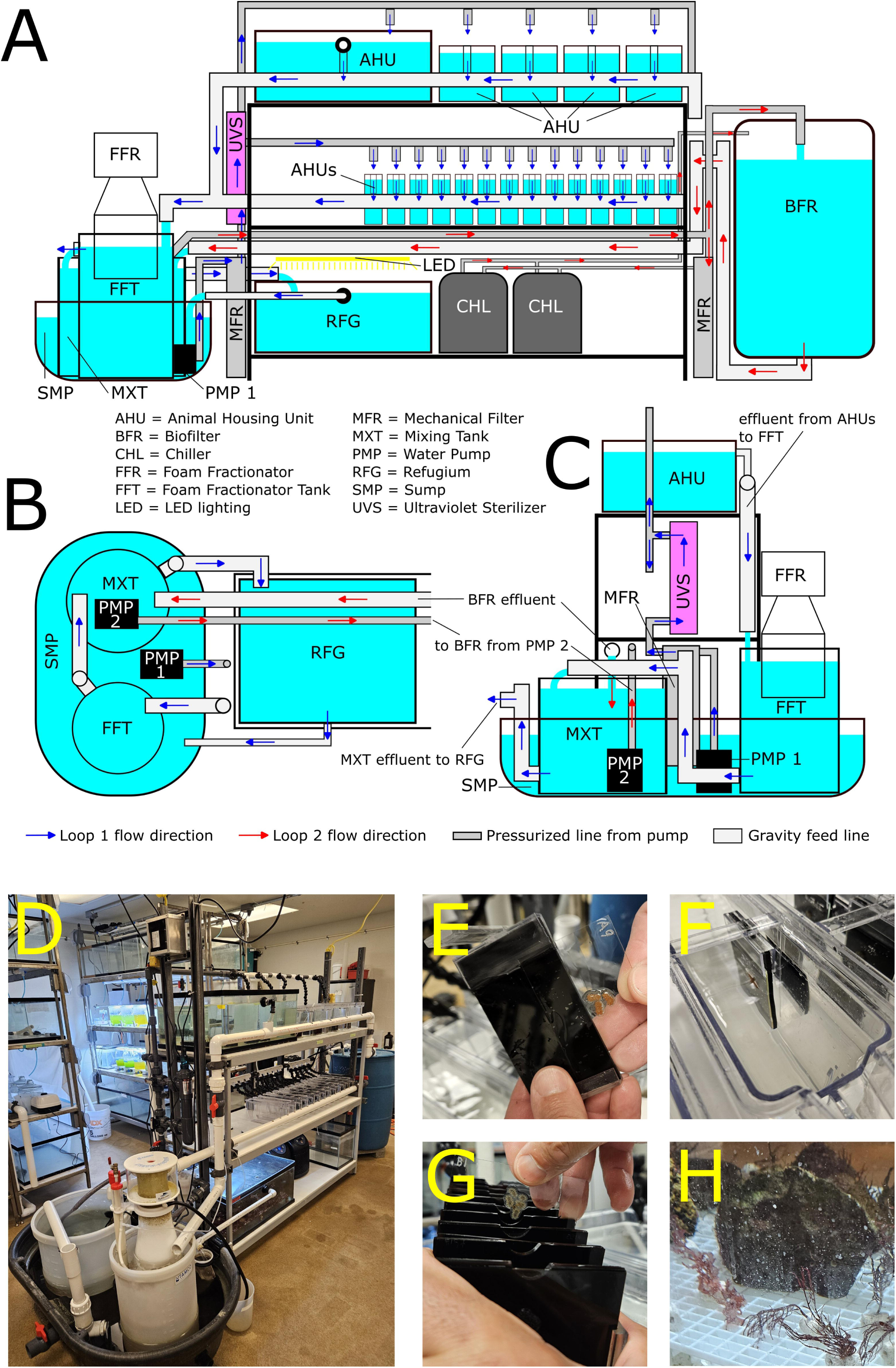
Completed revised RAS system. Schematic of revised RAS system with flow directionality in side view (A), top view of flow directionality between SMP components, RFG, and BFR (B), and view from SMP end (C). Images of black opaque backed slide holder system including single slide holders (E&F) and ten slide holder rack (G). Image of mature refugium containing two actively growing macrophyte species (H).

### 3.5. Revised RAS system growth

The few remaining B. schlosseri genotypes from the initial RAS design immediately began to recover in the newly optimized, revised RAS. They were able to fight off biofilm contamination and parasites, and systems increased in zooid number (Fig. 4A). In the optimized RAS, greater than 34 oozooids survived beyond 150 days from the first field collection after the RAS remodel (BM) with a mean RCZN of 16.5 over that time period in the selected genotypes.

**Figure 4.**
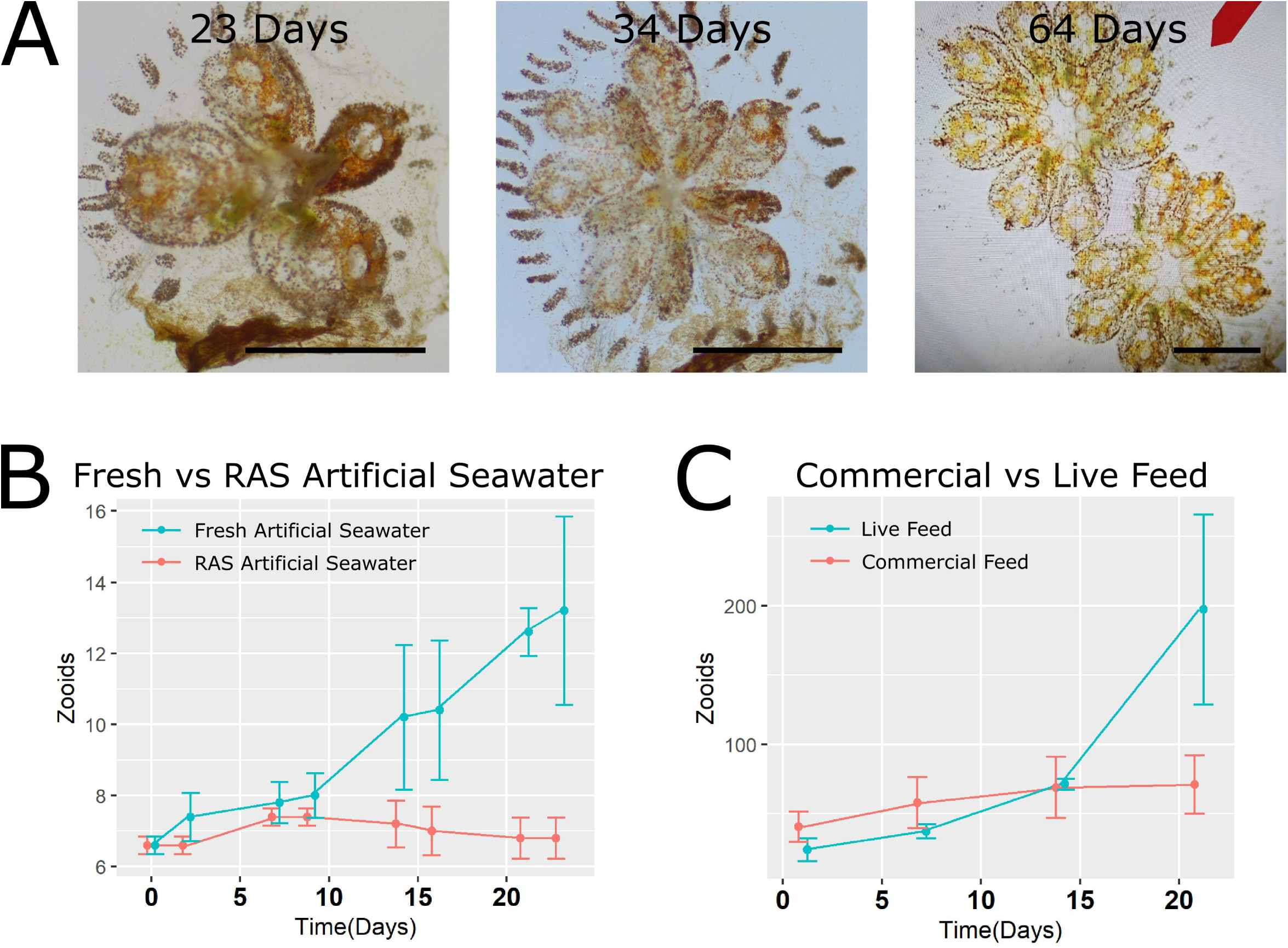
Growth of B. schlosseri in the optimized, revised RAS system and results of optimization experiments. Images showing recovery of a surviving genotype after transfer from the initial RAS system for the specified number of days to the optimized RAS system (A). Comparisons of B. schlosseri growth quantified by number of zooids/time(days) in RAS water versus fresh artificial seawater (AS) (B) and fed either live microalgae or commercial diets (C). Scale bars in each micrograph = 2 mm.

### 3.6. Superior growth of RAS adapted tunicates in static systems when using RAS water

When transferring tunicates from the optimized RAS to the static system a significant difference in growth rate was observed depending on whether they were grown using RAS water or freshly prepared AS. After a ∼10 day lag period, exponential growth was observed in the RAS water treated group while a reduction in zooid counts was observed in the fresh AS treated group (Fig 4B). The RAS treatment group yielded a mean RCZN of 1.95 that was significantly greater than the fresh AS group RCZN of 1.04 (p-value = 0.04952)

### 3.7. Elevated growth in response to a live feed (LF) mixture over commercial feed

After a ∼14 day lag period, an increase of colony growth rate was observed in LF-fed tunicates (Fig. 4C). The LF treatment group yielded a mean RCZN of 8.37 that was significantly greater than the CF group RCZN of 1.70 (p-value = 0.005587).

### 3.8. Feeding frequency increase in the optimized RAS system

An overall increased growth rate was apparent in multiple genotypes after increasing the feeding frequency from ever other day to daily with a rapid increase in zooid number and expansion into mutli-system colonies (Fig. 5A). This observation was confirmed by plotting of an additional zooid count 50 days post feeding frequency increase resulting in steeper slopes of zooid number/day for each genotype compared to the growth rates before increasing feeding frequency in the optimized RAS (Fig. 5B).

**Figure 5.**
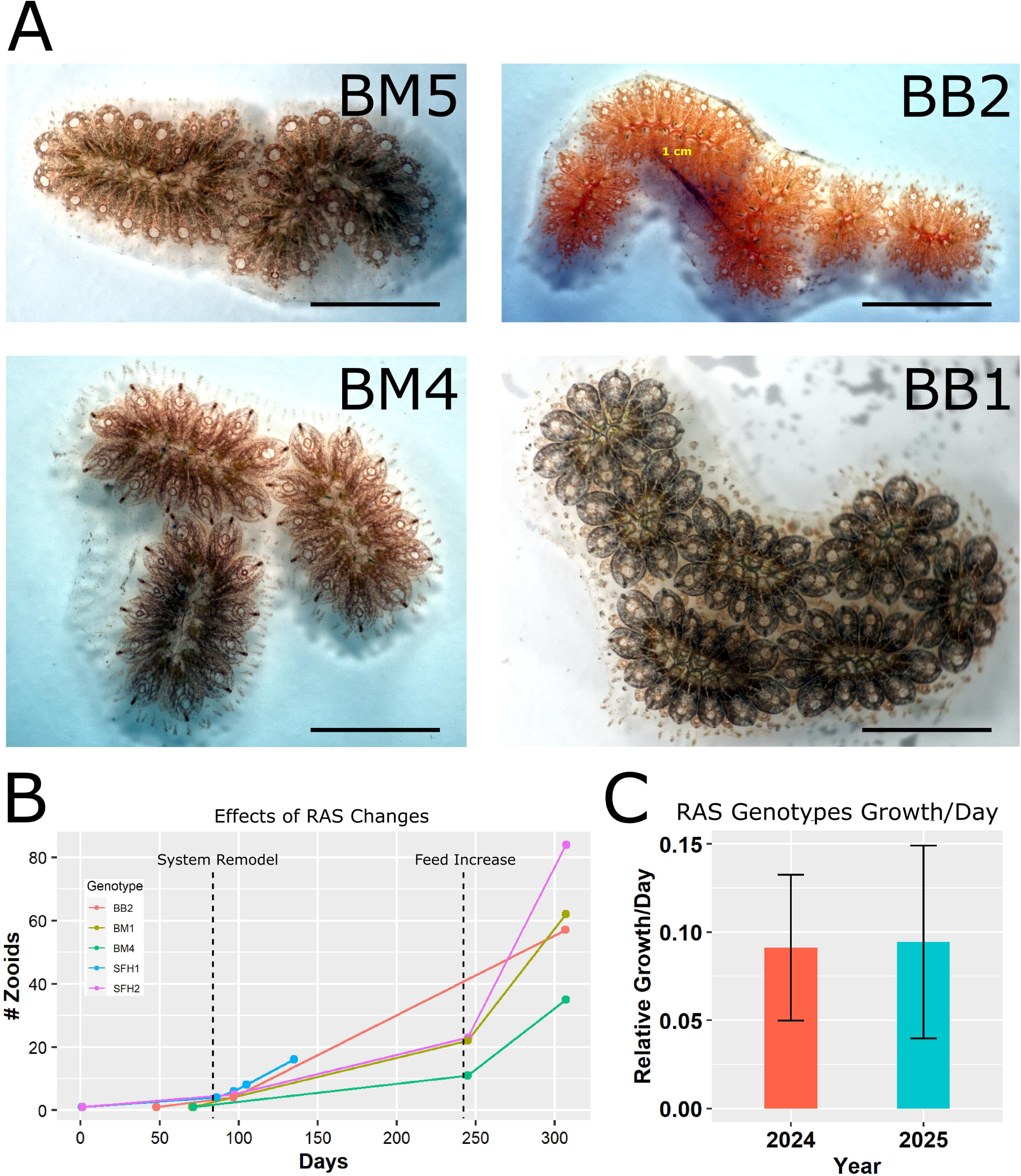
Representative images of B. schlosseri colonies from multiple genotypes after sustained growth in the optimized RAS system (A). Tracking of genotype growth (those for which there was zooid count data prior to RAS system remodel) over time with annotated time points of system remodel and feeding procedural changes (B). Growth rate (#systems/day) comparison of the same genotypes in 2024 and 2025 showing long-term maintenance of growth rates (C). Scale bars in each micrograph = 5 mm.

### 3.9. Long term status of B. schlosseri cultures in the optimized RAS

At the last 2024 sampling point, 34 genotypes remained with an average zooid count of 61 zooids/genotype. Currently, 11 genotypes have been maintained and asexually propagated for over 2 years with an average of 6 slides per genotype (∼6 systems/slide and ∼10 zooids/system) despite regular harvesting of animals for experiments (in 2025 number of systems per genotype were tracked to simplify record keeping). For example, 219 systems from genotype BM4 consisting of more than 6 zooids/per system were harvested for use in a study of copper toxicity stress[22]. Growth rate comparison of the current genotypes between 2024 and 2025 shows a maintenance of consistent growth rates over time (Fig. 5C). Additional signs of animal vigor in the optimized RAS system include production of gametes by many genotypes with a common occurrence of sexually produced oozooids on slides. This observation demonstrates the capacity of the optimized RAS for producing mixed genotypes in a controlled manner and to sustain a laboratory population by occasional sexual reproduction without additional sampling from the wild.

## 4. Discussion

The overall objective of this work was to construct a tunicate culture systems capable of sustaining and propagating healthy Botryllus schlosseri lab strains at a rate sufficient to support extensive research using this model organism. The static system resulting from these efforts was successful at achieving this goal., However, static system culture of tunicates has limited capacity at economic scale and an additional objective was to develop a higher capacity RAS system to: 1. reduce maintenance and cost of consumable resources; 2. create a more natural, biologically relevant culture environment; and 3. advance ecologically sustainable recirculating aquaculture system (RAS) techniques for these animals. An artificial culture system must balance replication of specific required environmental parameters, practical application and feasibility, and sufficient production output of animals to meet the goals of producing healthy animals for research at proper scale. The initial RAS version contained all the typical components of a closed loop marine culture system and maintained basic water quality parameter within acceptable limits for the culture of many marine invertebrates. Despite the apparent proper function of this initial RAS version, no B. schlosseri systems were able to grow larger than 6 zooids before completely regressing. This observation was in stark contrast to the exponential growth observed in the static system at this time.

### 4.1. Potential factors influencing B. schlosseri growth in closed system cultures

Many factors can potentially influence tunicate growth including abiotic physical parameters (temperature, salinity, lighting), density, food quality and availability, the presence or absence of inhibitory biofilms, pathogens, and interaction with allelopathic compounds. The abiotic parameters of the RAS system were consistent with those in the static system and standard conditions from the literature and were thus assumed not to be the primary reason for the differences in B. schlosseri growth. Intriguingly, in contrast to B. schlosseri, Botrylloides spp. grew steadily in the initial RAS version. Therefore, feed quantity was also not considered a primary factor for lack of sustained B. schlosseri growth, Rather, the presence of one or more other limiting factors specific to the interaction between B. schlosseri and the RAS initial version must be considered to explain the lack of sustained growth. In a closed loop system like RAS, the potential exist for accumulation of specific compounds if not effectively removed by the water treatment system. The weekly 100% water change of the static system would prevent any accumulation and consequences from long term retention of these wastes and harmful byproducts. With respect to many solutes, marine species are typically adapted to dilute environments and thus have low tolerance to toxins such as nitrogenous waste[23,24]. In an artificial system a dilute environment is relatively easy to achieve with low animal densities and only maintenance level feed. However, the goal of a production-focused system is to achieve high animal densities and rapid growth, which requires an abundant nutrition source and results in elevated waste. In a closed loop system, this can accelerate accumulation of inhibitory compounds including metabolic byproducts (such as nitrogenous waste) and allelopathic toxins from both target species, other species (Botrylloides spp.), and microbial decomposition of organics.

Nevertheless, many marine invertebrates, such as corals, can be proficiently cultured in RAS systems. Corals through their symbiont photosynthetic microbes, have the potential to simultaneously reduce potentially toxic nitrogenous waste in the ambient environment and convert them to nutrients for supporting growth[25]. Along with decomposing bacteria, they can assimilate and convert inorganic nitrogen back to useable nutrients[26]. In this regard corals would have an advantage over purely heterotrophic filter feeders (such as bivalve mollusks and tunicates) in closed loop recirculating systems which would require a higher density of suspended food to achieve rapid growth rates at the expense of water quality. Bivalves have been successfully cultured in RAS[17] but these animals have been shown to have higher tolerance to water quality parameters such as nitrogenous waste than other marine invertebrates[27]. From this study and others it is evident that at least some Botrylloides species are also amenable to RAS culture[19]. It is possible that B. schlosseri has a higher sensitivity to water quality than Botrylloides spp., rendering it more susceptible to nitrogenous waste toxicity below the detection limits of the methods used in this study or to some other metabolic waste byproduct. Excess waste can also negatively impact water quality indirectly through it’s influence on microbial communities. Excess inorganic nutrients and organics can support photosynthetic (i.e. cyanobacteria) and heterotrophic (i.e. Vibrio sp) harmful microbes, respectively, both of which are known to release exotoxins that are highly toxic to aquatic animals[28,29].

Allelopathic compounds generated by the tunicates themselves represent another potential factor for limiting B. schlosseri growth. The ability to inhibit establishment of nearby competitors of sessile marine invertebrates is a common phenomenon seen in sponges[30], corals[31] and anemones[32]. Many reports exist supporting a soluble factor released by colonial tunicates capable of suppressing growth of other tunicates species or genotypes[33,34]. Growth of Botrylloides leachii was observed to be roughly half in the proximity of other sessile invertebrates without physical contact relative to B. leachii grown in isolation[35]. These data indicate the presence of inhibitory soluble factors derived from competing organisms. Some of these ascidian-derived allelopathic compounds have been identified[36]. Multiple ascidian secreted alkaloids have demonstrated suppressive effects on general eukaryotic cell viability and proliferation[37–39]. It is possible that such competitive inhibitory factors are released by the tunicates and not efficiently removed by the filter system and accumulate in the circulating water. If such soluble compounds possess potent enough activity to have biological effects in an environment as dilute as the ocean, it is conceivable they could accumulate to impactful concentrations in a recirculating system.

### 4.2. RAS System Optimization

In an attempt to improve RAS of B. schlosseri, the system was remodeled to account for these potential deleterious factors discussed above. In the new configuration, foam fractionation was confined to an isolated chamber that received effluent directly from the AHUs from which the effluent flows to the biological filtration loop followed by the refugium. Foam fractionation is an effective treatment for purification of water[40] through removal of suspended solids[41] and dissolved organic carbon[42,43] including a variety of biochemically/pharmaceutically active compounds[44]. This arrangement allows for the most efficient removal of excess feed and deleterious tunicate byproducts, increases the retention time of the AHU effluent with the FF, and prevents mixing of AHU waste effluent with water destined to AHU distribution before passing through the complete water treatment cycle. This preliminary FF treatment of waste effluent can also improve BF performance as biological filter nitrification is more efficient with lower organic carbon loads[45].

This optimized arrangement also prevents the foam fractionator from removing potentially beneficial products generated by the refugium before it has the chance for distribution to the tunicates. Macrophytes release substantial dissolved and particulate organic matter into the ambient water[46,47], which is an additional potential nutrient source for the filter feeding tunicates. Macroalgae can improve water quality by assimilation of biological filtration end product inorganic nutrients[48] that can have deleterious effects on marine invertebrate physiology even at sublethal, low levels[23,49–51]. This removal of excess inorganic nutrients by macroalgae can also improve BF nitrification performance[52]. Additionally, macroalgae emit compounds that inhibit biofouling by microorganisms[53,54]. B. schlosseri and other tunicates, which are strongly inhibited by biofilms[55], often live in close association with macroalgae, and may receive benefits from these compounds. Such mutually beneficial interactions could explain why B. schlosseri typically require significant assistance in fighting biofilms by physical cleaning in laboratory conditions relative to the wild where they thrive amidst a complex biome.

Such symbiotic relationships with their natural environment may be partially replicated by a refugium housing macrophytes and substrates collected from the tunicate sampling sites and contribute to the improved B. schlosseri vigor in the optimized RAS system. Over the multiple years of this work, qualitative observation indicates a correlation between refugium and tunicate health. During a brief period when increased quantities of decaying organic matter were present (ie dying macrophytes), then the tunicate growth rate decreased. In contrast, when the refugium recovered and predominantly positive macrophyte growth was present, the condition and growth of the tunicates improved accordingly. The initial stages of refugium establishment were marked by a die-off of most macrophyte species introduced from the wild that were unable to adapt to the domesticated RAS conditions, followed by a period of establishment of a couple well adapted species that prevailed in a stable refugium. Thus, this maturation of the refugium represents a critical contribution toward the overall success of the optimized RAS system.

The establishment of a stable refugium was accompanied by dividing the flow between two pumps, one driving flow to the AHUs and another driving flow through a mechanical filter, the temperature control, and the biofilter. This arrangement allowed minimizing flow to the AHUs to increase retention time with the feed after feeding while maintaining high flows through the rest of the filtration system. Another notable change that contributed to the success of the optimized RAS version was holding tunicate containing glass slides in slide holders having a black, opaque back. This change was aimed at creating a more natural substrate for tunicate growth by preventing light exposure to the back of the slides and, thus, reducing competition with unwanted growth of photosynthetic microbes underneath the tunic.

### 4.3. Feed

The optimized fluidics of the RAS system, the maturation of the refugium, and removal of competitor Botrylloides spp. From the system all contributed to overall improved water quality and promoted long term survival, better condition, and moderate growth rates of B. schlosseri. However, the results from the static system clearly indicated that much higher growth rates were possible. A challenge of a production-focused RAS systems for a more sensitive species is maintenance of sufficient water quality despite high organic loads. Thus, it is critical to discern the ideal compromise between cleanliness of the water and feed availability for the target species. Feeding was initially minimized due to concerns of fouling the system with excess feed. However, comparison of optimized RAS system water with fresh AS from the static system indicated that the RAS water was as supportive of B. schlosseri growth as AS and the differences in growth rates between the two systems could be due flow regimen caused shorter feed retention in RAS compared to the static system.

Retention time with the feed and thus nutrient availability is a clear advantage of the static system and thus it was concluded that once whatever inhibitory factors were removed at the time of system remodel, then feed availability became the new limiting factor and the primary cause of the differences in growth rates observed between the static and revised RAS systems. This rationalization inspired the change of the RAS system feeding frequency from every other day to daily. Improved efficiency of excess feed removal from the system after optimization of the RAS flow regimen and water circuit allows for this increase in the amount of food without compromising water quality. The feed comparison experiment indicated that a live feed combination supported superior growth rates of B. schlosseri when compared to the commercial concentrate tested. An additional advantage of live feed is it’s motility and resistance to decay, in contrast to non-living commercial feed which can lead to non-beneficial microbial growth and contamination of the tunic. These attributes are especially important in long-term, continuous RAS culture where build-up of non-living organic matter can have negative consequences on water quality.

### 4.4. Future directions

Many of the key aspects influencing B. schlosseri culture were addressed during the course of this study, and proficient propagation of tunicates has been achieved using both static and RAS approaches. However, as total tunicate biomass increases and for further scaling up the capacity of RAS, it is possible the previous limiting factors could overwhelm the current system, requiring additional system improvements to sustain the same growth rates. Adding ozone (O_3_) to the FFT is an option that could increase the carrying capacity of the RAS system even further. As a powerful oxidant, ozone could inactive biochemical inhibitors, reduce the bioavailability of dissolved organics, and sterilize nuisance microorganisms. As the system is currently constructed, there is significant residence time in the treatment loop to dissipate O_3_ and UV exposure to revert residual O_3_ back to O_2_ before treated water returns to the animals. Additionally, the results from this work indicate animal retention time with the feed as a critical determinant of B. schlosseri growth and, thus, development of an automatic feeding system coupled with intermittently paused flow is worth considering if increased production capacity is needed.

## 5. Conclusions

Both systems described here have been demonstrated to be capable of long-term culture of healthy B. schlosseri over at least several years, each with specific strengths that can be used in combination to support research using this model organism. The static method requires more maintenance per animal/AHU but allows for isolation of animals into distinct units in which different experimental treatments can be applied. The RAS system does not allow for complete isolation of animal groups (e.g. different water quality and temperature) but can proficiently maintain and propagate separate genotype lines of B. schlosseri and serve as a source of laboratory strain animals for experiments. This workflow has several advantages over natural seawater-sourced systems as follows: As reproducible artificial systems, comparability of results is increased by eliminating variability associated with the source water, which can vary seasonally and stochastically. Other important advantages are increased biosecurity and containment of transgenic animals without the of risk genetically modified material escaping in the effluent to the environment. The cumulative modifications introduced during optimization of the RAS system and associated husbandry protocols guided by careful consideration of the most plausible influential factors has led to a long-term production capability of healthy animals sufficient to support extensive research efforts for over two years to date. Therefore, the systems documented here represent valuable resources for the scientific community to propel research using these intriguing tunicate models.

## CRediT authorship contribution statement

Jens Hamar: Writing – original draft, Visualization, Validation, Methodology, Investigation, Formal analysis, Conceptualization. Weizhen Dong: Conceptualization, Investigation. Brenda Luu: Conceptualization, Investigation. Mandy Lin: Investigation. Isabel Enriquez: Investigation. Maxime Leprêtre: Investigation. Alison M. Gardell: Funding acquisition, Conceptualization. Baruch Rinkevich: Funding acquisition, Conceptualization. Dietmar Kültz: Writing – review & editing, Supervision, Resources, Project administration, Funding acquisition, Conceptualization.

## Declaration of Competing Interest

The authors declare that they have no known competing financial interests or personal relationships that could have appeared to influence the work reported in this paper.

## Data availability

Data will be made available on request.

## Acknowledgements

This study was funded by National Science Foundation Grant MCB-2127516.

## Notes

### Competing Interest Statement

The authors have declared no competing interest.

## References

1. Carlson EA. How fruit flies came to launch the chromosome theory of heredity. Mutation Research/Reviews in Mutation Research. 2013 Jul 1;753(1):1–6.

2. Haffter P, Granato M, Brand M, Mullins MC, Hammerschmidt M, Kane DA, et al. The identification of genes with unique and essential functions in the development of the zebrafish, Danio rerio. Development. 1996 Dec 1;123(1):1–36.

3. Lew DJ, Dulić V, Reed SI. Isolation of three novel human cyclins by rescue of G1 cyclin (cln) function in yeast. Cell. 1991 Sep 20;66(6):1197–206.

4. Delsuc F, Philippe H, Tsagkogeorga G, Simion P, Tilak MK, Turon X, et al. A phylogenomic framework and timescale for comparative studies of tunicates. BMC Biology. 2018 Apr 13;16(1):39.

5. Taketa DA, De Tomaso AW. Botryllus schlosseri allorecognition: tackling the enigma. Dev Comp Immunol. 2015 Jan;48(1):254–65.

6. Rinkevich B. The colonial urochordate Botryllus schlosseri: from stem cells and natural tissue transplantation to issues in evolutionary ecology. Bioessays. 2002 Aug;24(8):730–40.

7. Voskoboynik A, Weissman IL. Botryllus schlosseri, an emerging model for the study of aging, stem cells, and mechanisms of regeneration. Invertebr Reprod Dev. 2015 Jan 30;59(sup1):33–8.

8. Gregorin C, Albarano L, Somma E, Costantini M, Zupo V. Assessing the Ecotoxicity of Copper and Polycyclic Aromatic Hydrocarbons: Comparison of Effects on Paracentrotus lividus and Botryllus schlosseri, as Alternative Bioassay Methods. Water. 2021;13(5).

9. Dunham (Epelbaum) A, Therriault T, Paulson A, Pearce C. Botryllid tunicates: Culture techniques and experimental procedures. Aquatic Invasions. 2009 Jan 1;4:111–20.

10. Liao PB, Mayo RD. Intensified fish culture combining water reconditioning with pollution abatement. Aquaculture. 1974 Feb 1;3(1):61–85.

11. Heinen JM, Hankins JA, Adler PR. Water quality and waste production in a recirculating trout-culture system with feeding of a higher-energy or a lower-energy diet. Aquaculture Research. 1996 Sep 1;27(9):699–710.

12. Khanjani MH, Sharifinia M, Hajirezaee S. Strategies for promoting sustainable aquaculture in arid and semi-arid areas – A review. Annals of Animal Science. 2024;24(2):293–305.

13. Gopakumar G, George RM, Jasmine S. Breeding and larval rearing of the clownfish Amphiprion chrysogaster. In 1999. Available from: https://api.semanticscholar.org/CorpusID:82863443

14. Tartila S. The Clownfish (Amphiprion spp.) Larviculture Technique with Recirculating Aquaculture System (RAS) in Buleleng, Bali. Journal of Aquaculture Development and Environment. 2023 Mar 1;6:363–9.

15. Weirich CR, Riley KL, Riche M, Main KL, Wills PS, Illán G, et al. The status of Florida pompano, , as a commercially ready species for U.S. marine aquaculture. Journal of the World Aquaculture Society. 2021 Jun 1;52(3):731–63.

16. Grosjean P, Spirlet C, Gosselin P, Vaitilingon D, Jangoux M. Land-based, closed-cycle echiniculture of Paracentrotus lividus (Lamarck) (Echinoidea: Echinodermata): A long-term experiment at a pilot scale. Journal of Shellfish Research. 1998 Dec 1;17:1523–31.

17. Kuhn DD, Angier MW, Barbour SL, Smith SA, Flick GJ. Culture feasibility of eastern oysters (Crassostrea virginica) in zero-water exchange recirculating aquaculture systems using synthetically derived seawater and live feeds. Aquacultural Engineering. 2013 May 1;54:45–8.

18. Lunden JJ, Turner JM, McNicholl CG, Glynn CK, Cordes EE. Design, development, and implementation of recirculating aquaria for maintenance and experimentation of deep-sea corals and associated fauna. Limnology and Oceanography: Methods. 2014 Jun 1;12(6):363–72.

19. Wawrzyniak MK, Matas Serrato LA, Blanchoud S. Artificial seawater based long-term culture of colonial ascidians. Developmental Biology. 2021 Dec 1;480:91–104.

20. Rinkevich B, Shapira M. An improved diet for inland broodstock and the establishment of an inbred line from Botryllus schlosseri, a colonial sea squirt (Ascidiacea). Aquatic Living Resources. 1998 May 1;11(3):163–71.

21. Taketa DA, Nydam ML, Langenbacher AD, Rodriguez D, Sanders E, De Tomaso AW. Molecular evolution and in vitro characterization of Botryllus histocompatibility factor. Immunogenetics. 2015 Oct 1;67(10):605–23.

22. Leprêtre M, Kültz D. Copper-Induced Stress and Recovery Impacts on Organismal Phenotypes and the Underlying Proteomic Signatures in <em>Botryllus schlosseri</em>. bioRxiv. 2025 Jan 1;2025.06.12.659394.

23. Muir PR, Sutton DC, Owens L. Nitrate toxicity toPenaeus monodon protozoea. Marine Biology. 1991 Feb 1;108(1):67–71.

24. Phillips BM, Nicely PA, Hunt JW, Anderson BS, Tjeerdema RS, Palmer FH. Tolerance of Five West Coast Marine Toxicity Test Organisms to Ammonia. Bulletin of Environmental Contamination and Toxicology. 2005 Jul 1;75(1):23–7.

25. Leal MC, Ferrier-Pagès C, Petersen D, Osinga R. Coral aquaculture: applying scientific knowledge to ex situ production. Reviews in Aquaculture. 2016 Jun 1;8(2):136–53.

26. Pernice M, Meibom A, Van Den Heuvel A, Kopp C, Domart-Coulon I, Hoegh-Guldberg O, et al. A single-cell view of ammonium assimilation in coral-dinoflagellate symbiosis. ISME J. 2012 Jul;6(7):1314–24.

27. Epifanio CE, Srna RF. Toxicity of ammonia, nitrite ion, nitrate ion, and orthophosphate to Mercenaria mercenaria and Crassostrea virginica. Marine Biology. 1975 Dec 1;33(3):241–6.

28. Vandeputte M, Kashem MdA, Bossier P, Vanrompay D. Vibrio pathogens and their toxins in aquaculture: A comprehensive review. Reviews in Aquaculture. 2024 Sep 1;16(4):1858–78.

29. Zanchett G, Oliveira-Filho EC. Cyanobacteria and cyanotoxins: from impacts on aquatic ecosystems and human health to anticarcinogenic effects. Toxins (Basel). 2013 Oct 23;5(10):1896–917.

30. Thompson JE. Exudation of biologically-active metabolites in the sponge Aplysina fistularis. Marine Biology. 1985 Aug 1;88(1):23–6.

31. Maida M, Sammarco PW, Coll JC. Effects of soft corals on scleractinian coral recruitment. I: Directional allelopathy and inhibition of settlement. Mar Ecol Prog Ser. 1995;121:191–202.

32. Bak RPM, Borsboom JLA. Allelopathic interaction between a reef coelenterate and benthic algae. Oecologia. 1984 Aug 1;63(2):194–8.

33. Paetzold C, Giberson DJ, Hill JV, Davidson JDP, Davidson J. Effect of colonial tunicate presence on Ciona intestinalis recruitment within a mussel farming environment. Management of Biological Invasions. 2012;3:15–23.

34. Young CM, Chia FS. LABORATORY EVIDENCE FOR DELAY OF LARVAL SETTLEMENT IN RESPONSE TO A DOMINANT COMPETITOR. International Journal of Invertebrate Reproduction. 1981 Jan 1;3(4):221–6.

35. Myers PE. Space versus other limiting resources for a colonial tunicate, Botrylloides leachii (Savigny), on fouling plates. Journal of Experimental Marine Biology and Ecology. 1990 Aug 21;141(1):47–52.

36. Ramesh C, Tulasi BR, Raju M, Thakur N, Dufossé L. Marine Natural Products from Tunicates and Their Associated Microbes. Mar Drugs. 2021 May 26;19(6).

37. Roberge M, Berlinck RGS, Xu L, Anderson HJ, Lim LY, Curman D, et al. High-Throughput Assay for G2 Checkpoint Inhibitors and Identification of the Structurally Novel Compound Isogranulatimide1. Cancer Research. 1998 Dec 1;58(24):5701– 6.

38. Batista PJ, Nuzzo G, Gallo C, Carbone D, dell’Isola M, Affuso M, et al. Chemical and Pharmacological Prospection of the Ascidian Cystodytes dellechiajei. Marine Drugs. 2024;22(2).

39. Abourriche A, Abboud Y, Maoufoud S, Mohou H, Seffaj T, Charrouf M, et al. Cynthichlorine: a bioactive alkaloid from the tunicate Cynthia savignyi. Il Farmaco. 2003 Dec 1;58(12):1351–4.

40. Buckley T, Xu X, Rudolph V, Firouzi M, Shukla P. Review of foam fractionation as a water treatment technology. Separation Science and Technology. 2021 Jul 18;57:1–30.

41. Weeks NC, Timmons MB, Chen S. Feasibility of using foam fractionation for the removal of dissolved and suspended solids from fish culture water. Aquacultural Engineering. 1992 Jan 1;11(4):251–65.

42. Roselet M, Roselet F, Abreu PC. Foam fractionator as a tool to remove dissolved organic matter and improve the flocculation of the marine microalga Nannochloropsis oceanica. Journal of Applied Phycology. 2019 Oct 1;31(5):2911– 9.

43. von Ahnen M, Stedmon CA, Hambly AC. Removal of dissolved organic matter from the woodchip bioreactor start-up by foam fractionation. Water Science and Technology. 2023 Mar 14;87(6):1454–64.

44. Datta P, Ghosh A, Chakraborty P, Gangopadhyay A, Prabir M, Datta K. Foam Fractionation in Separation of Pharmaceutical Biomolecules: A Promising Unit Operation for Industrial Process and Waste Control Introduction The recovery, isolation and purification of biomolecules like pharmaceutical proteins in pharmaceutical. JOURNAL OF PHARMACEUTICAL AND FUNDAMENTAL RESEARCH. 2015 Sep 26;20153:33–41.

45. Michaud L, Blancheton JP, Bruni V, Piedrahita R. Effect of particulate organic carbon on heterotrophic bacterial populations and nitrification efficiency in biological filters. Aquacultural Engineering. 2006 May 1;34(3):224–33.

46. Haas AF, Naumann MS, Struck U, Mayr C, el-Zibdah M, Wild C. Organic matter release by coral reef associated benthic algae in the Northern Red Sea. Journal of Experimental Marine Biology and Ecology. 2010 Jun 30;389(1):53–60.

47. Hauri C, Fabricius KE, Schaffelke B, Humphrey C. Chemical and physical environmental conditions underneath mat- and canopy-forming macroalgae, and their effects on understorey corals. PLoS One. 2010 Sep 13;5(9):e12685.

48. Copertino M da S, Tormena T, Seeliger U. Biofiltering efficiency, uptake and assimilation rates of Ulva clathrata (Roth) J. Agardh (Clorophyceae) cultivated in shrimp aquaculture waste water. Journal of Applied Phycology. 2009 Feb 1;21(1):31–45.

49. Böttger SA, McClintock JB. Effects of inorganic and organic phosphate exposure on aspects of reproduction in the common sea urchin Lytechinus variegatus (Echinodermata: Echinoidea). Journal of Experimental Zoology. 2002 Jun 1;292(7):660–71.

50. Zhu B, Wang G, Huang B, Tseng CK. Effects of temperature, hypoxia, ammonia and nitrate on the bleaching among three coral species. Chinese Science Bulletin. 2004 Sep 1;49(18):1923–8.

51. Marubini F, Davies PS. Nitrate increases zooxanthellae population density and reduces skeletogenesis in corals. Marine Biology. 1996 Dec 1;127(2):319–28.

52. Li X, Deng Y, Li X, Ma X, Wang J, Li J. Integration of Marine Macroalgae (Chaetomorpha maxima) with a Moving Bed Bioreactor for Nutrient Removal from Maricultural Wastewater. Archaea. 2020;2020:8848120.

53. Bhadury P, Wright PC. Exploitation of marine algae: biogenic compounds for potential antifouling applications. Planta. 2004 Aug 1;219(4):561–78.

54. Dahms HU, Dobretsov S. Antifouling Compounds from Marine Macroalgae. Marine Drugs. 2017;15(9).

55. Zapata M, Silva F, Luza Y, Wilkens M, Riquelme C. The inhibitory effect of biofilms produced by wild bacterial isolates to the larval settlement of the fouling ascidia Ciona intestinalis and Pyura praeputialis. Electronic Journal of Biotechnology. 2007 Jan;10:149–59.

